# Evaluating Metagenome Assembly on a Simple Defined Community with Many Strain Variants

**DOI:** 10.1101/155358

**Authors:** Sherine Awad, Luiz Irber, C. Titus Brown

## Abstract

We evaluate the performance of three metagenome assemblers, IDBA, MetaSPAdes, and MEGAHIT, on short-read sequencing of a defined “mock” community containing 64 genomes (Shakya et al. (2013)). We update the reference metagenome for this mock community and detect several additional genomes in the read data set. We show that strain confusion results in significant loss in assembly of reference genomes that are otherwise completely present in the read data set. In agreement with previous studies, we find that MEGAHIT performs best computationally; we also show that MEGAHIT tends to recover larger portions of the strain variants than the other assemblers.

## Introduction

Metagenomics refers to sequencing of DNA from a mixture of organisms, often from an environmental or uncultured sample. Unlike whole genome sequencing, metagenomics targets a mixture of genomes, which introduces metagenome-specific challenges in analysis [1]. Most approaches to analyzing metagenomic data rely on mapping or comparing sequencing reads to reference sequence collections. However, reference databases contain only a small subset of microbial diversity [2], and much of the remaining diversity is evolutionarily distant and reference-based search techniques may not recover it [3].

As sequencing capacity increases and sequence data is generated from many more environmental samples, metagenomics is increasingly using *de novo* assembly techniques to generate new reference genomes and metagenomes [4]. There are a number of metagenome assemblers that are widely used - see [5] for an overview of the available software, and [1] for a review of the different assembler methodologies. However, evaluating the results of these assemblers is challenging due to the general lack of good quality reference metagenomes.

Moya et al. in [6] evaluated metagenome assembly using two simulated 454 viral metagenome and six assemblers. The assemblies were evaluated based on several metrics including N50, percentages of reads assembled, accuracy when compared to the reference genome. In addition to these metrics, the authors evaluated, chimeras per contigs and the effect of assembly on taxonomic and functional annotations.

Mavromatis et al. in [7] provided a benchmark study to evaluate the fidelity of metagenome processing methods. The study used simulated metagenomic data sets constructed at different complexity levels. The datasets were assembled using Phrap v3.57, Arachne v.2 [8] and JAZZ [9]. This study evaluates assembly, gene prediction, and binning methods. However, the study did not evaluate the assembly quality against a reference genome.

Rangwala et al. in [10] presented an evaluation study of metagenome assembly. The study used a de Bruijn graph based assembler ABYSS [11] to assemble simulated metagenome reads of 36 bp. The data set is classified at different complexity levels. The study compared the quality of the assembly of the data sets in terms of contig length and assembly accuracy. The study also took into consideration the effect of kmer size and the degree of chimericity. However, the study evaluated the assembly based on only one assembler. Also, these previous studies used simulated data, which may lack confounders of assembly such as sequencing artifacts and GC bias.

In a landmark study, Shakya et al. (2013) constructed a synthetic community of organisms by mixing DNA isolated from individual cultures of 64 bacteria and archaea, including a variety of strains across a range of average nucleotide distances [12]. In addition to performing 16s amplicon analysis and doing 454 sequencing, the authors shotgun-sequenced the mixture with Illumina. While the authors concluded that this metagenomic sequencing generally outperformed amplicon sequencing, they did not conduct an assembly based analysis. This data set was also used in several other evaluation studies, including gbtools for binning [13] and benchmarking of the MEGAHIT assembler [14].

More recently, several benchmark studies systematically evaluated metagenome assembly of short reads. The Critical Assessment of Metagenome Interpretation (CAMI) collaboration benchmarked a number of metagenome assemblers on several data sets of varying complexity, evaluating recovery of novel genomes and multiple strain variants [3]. Notably, CAMI concluded that “The resolution of strain-level diversity represents a substantial challenge to all evaluated programs.” Another recent study evaluated eight assemblers on nine environmental metagenomes and three simulated data sets and provided a workflow for choosing a metagenome assembler based on the biological goal and computational resources available [15]. [5] explored metagenome assembler performance on a pair of real data sets, again concluding that the biological goal and computational resources defined the choice of assembler. Also see [16] for an analysis of a previously generated HMP benchmark data set; however, the Illumina reads used for this study are much shorter than current sequencing and are arguably not relevant to future studies.

In this study, we extend previous work by delving into questions of chimeric misassembly and strain recovery in the Shakya et al. (2013) data set. First, we update the list of reference genomes for Shakya et al. to include the latest GenBank assemblies along with plasmids. We then compare IDBA [17], MetaSPAdes [18], and MEGAHIT [19] performance on assembling this short-read data set, and explore concordance in recovery between the three assemblers. We describe the effects of “strain confusion” between multiple strains. We also detect and analyze several previously unreported strains and genomes in the Shakya et al. data set. We find that in the absence of closely related genomes, all three metagenome assemblers recover 95% or more of known reference genomes. However, in the presence of closely related genomes, these three metagenome assemblers vary widely in their performance and, in extreme cases, can fail to recover the majority of some genomes even when they are completely present in the reads. Our report provides strong guidance on choice of assemblers and extends previous analyses of this low-complexity metagenome benchmarking data set.

### Datasets

We used a diverse mock community data set constructed by pooling DNA from 64 species of bacteria and archaea and sequencing them with Illumina HiSeq. The raw data set consisted of 109,629,496 reads from Illumina HiSeq 101 bp paired-end sequencing (2x101) with an untrimmed total length of 11.07 Gbp and an estimated fragment size of 380 bp [12].

The original reads are available through the NCBI Sequence Read Archive at Accession SRX200676. We updated the 64 reference genomes sets from NCBI GenBank using the latest available assemblies with plasmid content (June 2017); the accession numbers are available as `accession-list-ref.txt` in the Zenodo repository, DOI: 10.5281/zenodo.821919. For convenience, the updated reference genome collection is available for download at the archival URL https://osf.io/vbhy5/.

## Methods

The analysis code and run scripts for this paper are written in Python and bash, and are available at https://github.com/dib-lab/2016-metagenome-assembly-eval/ (archived at Zenodo DOI: 10.5281/zenodo.821919). The scripts and overall pipeline were examined by the first and senior authors for correctness. In addition, the bespoke reference-based analysis scripts were tested by running them on a single-colony *E. coli* MG1655 data set with a high quality reference genome [20].

### Quality Filtering

We removed adapters with Trimmomatic v0.30 in paired-end mode with the TruSeq adapters [21], using light quality score trimming (`LEADING:2 TRAILING:2 SLIDINGWINDOW:4:2 MINLEN:25`) as recommended in MacManes, 2014 [22].

### Reference Coverage Profile

To evaluate how much of the reference metagenome was contained in the read data, we used `bwa aln` (v0.7.7.r441) to map reads to the reference enome [23]. We then calculated how many reference bases were covered by mapped reads (custom script `coverage-profile.py`).

### Measuring k-mer inclusion and Jaccard similarity

We used MinHashing as implemented in sourmash to estimate k-mer inclusion and Jaccard similarity between data sets [24]. MinHash signatures were prepared with `sourmash compute` using –`scaled 10000`. K-mer inclusion was computed by taking the ratio of the number of intersecting hashes with the query over the total number of hashes in the subject MinHash. Jaccard similarity was computed as in [25] by taking the ratio of the number of intersecting hashes between the query and subject over the number of hashes in the union. K-mer sizes for comparison were chosen at 21, 31, or 51, depending on the level of taxonomic specificity desired - genus, species, or strain, respectively, as described in [26].

Where specified, high-abundance k-mers were selected for counting by using the script `trim-low-abund.py` script with `-C 5` from khmer v2 [27, 28].

### Assemblers

We assembled the quality-filtered reads using three different assemblers: IDBA-UD [17], MetaSPAdes [18], and MEGAHIT [19]. For IDBA-UD v1.1.3 [17], we used –`pre_correction` to perform pre-correction before assembly and -r for the pe files. IDBA could not ingest orphan sequences so singleton reads were omitted from this assembly.

For MetaSPAdes v3.10.1 [18], we used –`meta –pe1-12 –pe1-s` where `#x2014;meta` is used for metagenomic data sets, –`pe1-12` specifies the interlaced reads for the first paired-end library, and `–pe1-s` provides the orphan reads remaining from quality trimming.

For MEGAHIT v1.1.1-2-g02102e1 [19], we used `-l 101 -m 3e9 –cpu-only` where `-l` is for maximum read length, `-m` is for max memory in bytes to be used in constructing the graph, and –`cpu-only` uses only the CPU and no GPUs. We also used –`presets meta-large` for large and complex metagenomes, and –`12` and `-r` to specify the interleaved-paired-end and single-end files respectively. MEGAHIT allows the specification of a memory limit and we used `-M 1e+10` for 10 GB.

All three assemblies were executed on the same XSEDE Jetstream instance (S1.Xxlarge) at Indiana University, running Ubuntu 16.04 (install 6/21/17, Ubuntu 16.04 LTS Development + GUI support + Docker; based on Ubuntu cloud image for 16.04 LTS with basic dev tools, GUI/Xfce added). Assemblers were limited to 16 threads. We recorded RAM and CPU time for each assembly using `/usr/bin/time -v`. Install and execute details as well as output timings and logs are available in the `pipeline/runstats` directory of the Zenodo archive.

Unless otherwise mentioned, we eliminated all contigs less than 500 bp from each assembly prior to further analysis.

### Mapping

We aligned all quality-filtered reads to the reference metagenome with bwa aln (v0.7.7.r441) [23]. We aligned paired-end and orphaned reads separately. We then used samtools (v0.1.19) [29] to convert SAM files to BAM files for both paired-end and orphaned reads. To count the unaligned reads, we included only those records with the “4” flag in the SAM files [29].

### Assembly analysis using NUCmer

We used the NUCmer tool from MUMmer3.23 [30] to align assemblies to the reference genome with options `–coords –p`. Then we parsed the generated “.coords” file using a custom script `analyze_assembly.py`, and calculated several analysis metrics across all three assemblies at a 99% alignment identity.

### Reference-based analysis of the assemblies

We conducted reference-based analysis of the assemblies under two conditions. “Loose” alignment conditions used all available alignments, including redundant and overlapping alignments. “Strict” alignment conditions took only the longest alignment for any given contig, eliminating all other alignments.

The script `summarize-coords2.py` was used to calculate aligned coverage from the loose alignment conditions: each base in the reference was marked as “covered” if it was included in at least one alignment. The script `analyze_ng50.py` was used to calculate NGA 50 for each individual reference genome.

### Analysis of chimeric misassemblies

We analyzed each assembly for chimeric misassemblies by counting the number of contigs that contained matches to two distinct reference genomes. In order to remove secondary alignments from consideration, we included only the longest non-overlapping NUCmer alignments for each contig at a minimum alignment identity of 99%. We then used the script `analyze_chimeric2.py` to find individual contigs that matched more than one distinct reference genome. As a negative control on our analysis, we verified that this approach yielded no positive results when applied to the alignments of the reference metagenome against itself.

### Analysis of unmapped reads

We conducted assembly and analysis of unmapped reads with MEGAHIT, NUCmer, and sourmash as above. The new GenBank genomes are listed in the Zenodo archive at the file `accession-list-unmapped.txt` and for convenience are available for download at the archival URL https://osf.io/34ef8/.

## Results

### The raw data is high quality

The reads contain 11,072,579,096 bp (11.07 Gbp) in 109,629,496 reads with 101.0 average length (2x101bp Illumina HiSeq).

Trimming removed 686,735 reads (0.63%). After trimming, we retained 108,422,358 paired reads containing 10.94 Gbp with an average length of 100.9 bases. A total of 46.56 Mbp remained in 520,403 orphan reads with an average length of 89.5 bases. In total, the quality trimmed data contained 10.98 Gbp in 108,942,761 reads. This quality trimmed (“QC”) data set was used as the basis for all further analyses.

### The reference metagenome is not completely present in the reads

We next evaluated the fraction of the reference genome covered by at least one read (see Methods for details). Quality filtered reads cover 203,058,414 (98.76%) bases of the reference metagenome (205,603,715 bp total size). Figure 1 shows the cumulative coverage profile of the reference metagenome, and the percentage of bases with that coverage. Most of the reference metagenome was covered at least minimally; only 3.33% of the reference metagenome had mapping coverage <5, and 1.24% of the bases in the reference were not covered by any reads in the QC data set.

**Figure 1:**
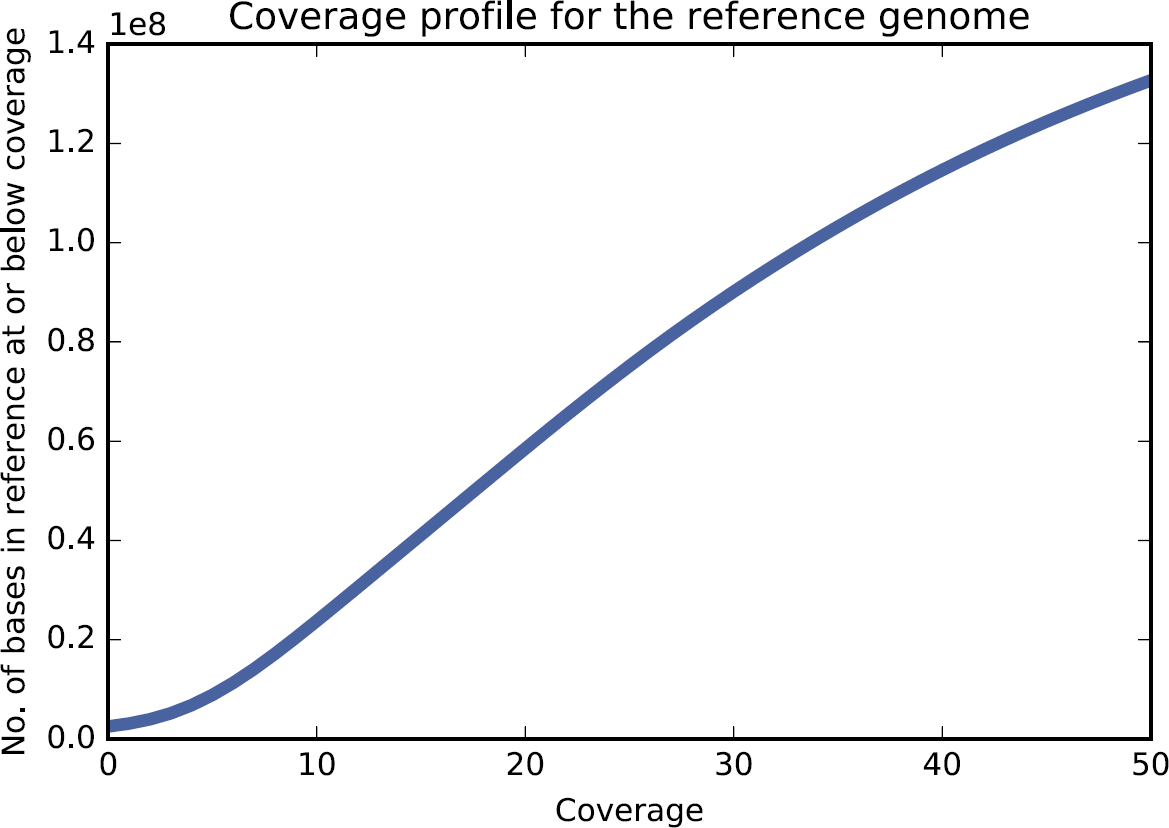
Cumulative coverage profile for the reference metagenome, based on read mapping.

In order to evaluate reconstructability with De Bruijn graph assemblers, we next examined k-mer containment of the reference in the reads for *k* of 21, 31, 41, and 51 (Table 1). The k-mer overlap decreases from 96.8% to 94.1% as the k-mer size increases. This could be caused by low coverage of some portions of the reference and/or variation between the reads and the reference.

**Table 1:** Jaccard containment of the reference in the reads

| k-mer size | % reference in reads |
| --- | --- |
| 21 | 96.8% |
| 31 | 95.9% |
| 41 | 94.9% |
| 51 | 94.1% |

### Some individual reference genomes are poorly represented in the reads

To see if specific reference genomes exhibited low coverage, we analyzed read mapping coverage for individual genomes. Of the 64 reference genomes used in the metagenome, 60 had a per-base mapping coverage above 95%. The remaining four varied significantly (Table 2), with *F. nucleatum* the lowest – only 47.6% of the bases in the reference genome are covered by one or more mapped reads.

**Table 2:**
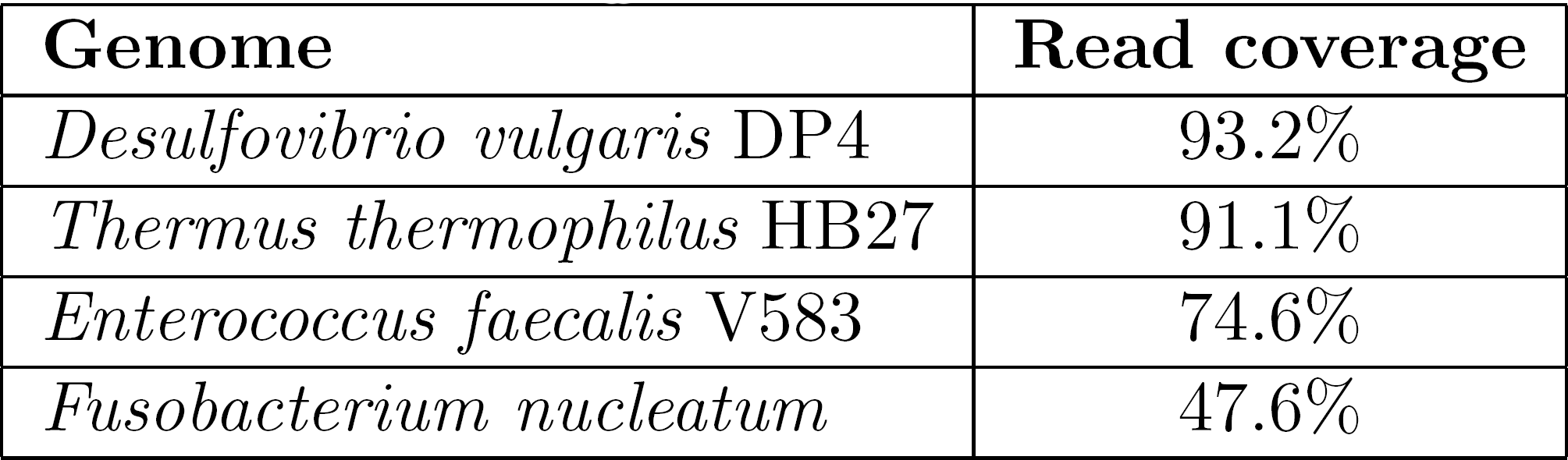
Top uncovered genomes

| Genome | Read coverage |
| --- | --- |
| *Desulfovibrio vulgaris* DP4 | 93.2% |
| *Thermus thermophilus* HB27 | 91.1% |
| *Enterococcus faecalis* V583 | 74.6% |
| *Fusobacterium nucleatum* | 47.6% |

We next did a 51-mer containment analysis of each reference genome in the reads; k=51 was chosen so as to be specific to strain content [26]. 99% or more of the constituent 51-mers for 51 of the 64 reference genomes were present in the reads, suggesting that each of the 51 genomes was entirely present at some minimal coverage.

We excluded the remaining 13 genomes (see Table 3) from any further reference-based analysis because interpreting recovery and misassembly statistics for these genomes would be confounding; also see the discussion of strain variants, below.

**Table 3:** Genomes removed from reference for low 51-mer presence

| 51-mers in reads | Genome |
| --- | --- |
| 98.7 | *Leptothrix cholodnii* |
| 98.7 | *Haloferax volcanii* DS2 |
| 98.6 | *Salinispora tropica* CNB-440 |
| 97.4 | *Deinococcus radiodurans* |
| 97.2 | *Zymomonas mobilis* |
| 97.1 | *Ruegeria pomeroyi* |
| 96.8 | *Shewanella baltica* OS223 |
| 95.5 | *B. bronchiseptica* D989 |
| 94.5 | *Burkholderia xenovorans* |
| 72.0 | *Desulfovibrio vulgaris* DP4 |
| 65.0 | *Thermus thermophilus* HB27 |
| 53.4 | *Enterococcus faecalis* |
| 4.7 | *Fusobacterium nucleatum* ATCC 25586 |

### MEGAHIT is the fastest and lowest-memory assembler evaluated

We ran three commonly used metagenome assemblers on the QC data set: IDBA-UD, MetaSPAdes, and MEGAHIT. We recorded the time and memory usage of each (Table 4). In computational requirements, MEGAHIT outperformed both MetaSPAdes and IDBA-UD, , producing an assembly in 1.5 hours (“wall time”) – 1.6 times faster than IDBA and 2.6 times faster than MetaSPAdes. MEGAHIT used only 10 GB of RAM as requested – about 60% of the memory used by IDBA and a third of the memory used by IDBA and MetaSPAdes, respectively. CPU time measurements (which include processing on multiple CPU cores) show that all three assemblers use multiple cores effectively.

**Table 4:**
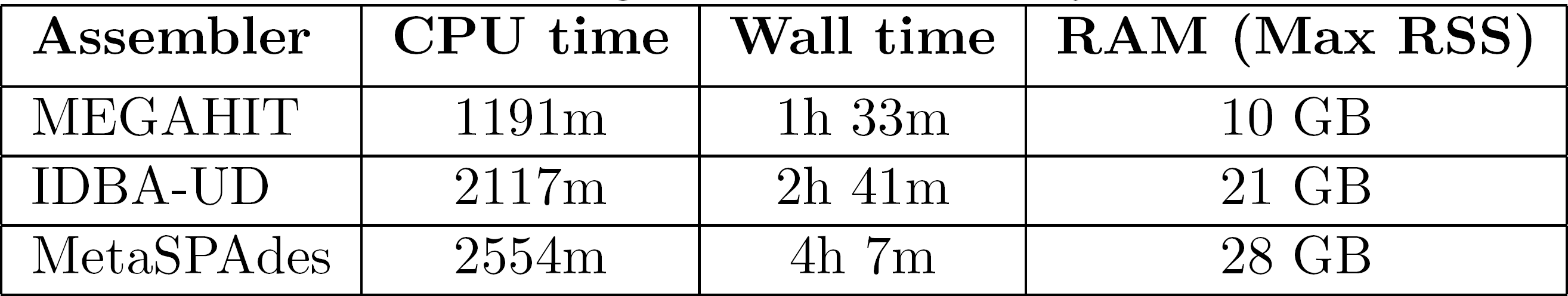
Running Time and Memory Utilization

### The assemblies contain most of the raw data

We assessed read inclusion in assemblies by mapping the QC reads to the length-filtered assemblies and counting the remaining unmapped reads. Depending on the assembly, between 2.7 million and 3.9 million reads (2.5-3.5%) did not map to the assemblies (Table 5). All of the assemblies included the large majority of high-abundance 51-mers (more than 96.8% in all cases).

**Table 5:** Read and high-abundance (> 5) k-mer exclusion from assemblies

| Assembly | Unmapped Reads | 51-mers omitted |
| --- | --- | --- |
| IDBA | 3,328,674 (3.05%) | 2.4% |
| MetaSPAdes | 3,844,123 (3.52%) | 3.2% |
| MEGAHIT | 2,737,640 (2.51%) | 2.8% |

### Much of the reference is covered by the assemblies

We next evaluated the extent to which the assembled contigs recovered the “known/true” metagenome sequence by aligning each assembly to the adjusted reference (Table 6). Each of the three assemblers generates contigs that cover more than 93.1% of the reference metagenome at high identity (99%) with little duplication (approximately 1%). All three assemblies contain between 96.2% and 97.2% of the 51-mers in the reference.

**Table 6:** Contig coverage of reference with loose alignment conditions.

| Assembly | bases aligned | duplication | 51-mers |
| --- | --- | --- | --- |
| MEGAHIT | 94.8% | 1.0% | 96.7% |
| MetaSPAdes | 93.1% | 1.1% | 96.2% |
| IDBA | 93.6% | 0.98% | 97.2% |

At 99% identity with the loose mapping approach, approximately 2.5% of the reference is missed by all three assemblers, while 1.7% is uniquely covered by MEGAHIT, 0.74% is uniquely covered by MetaSPAdes, and 0.64% is uniquely covered by IDBA.

### The generated contigs are broadly accurate

When counting only the best (longest) alignment per contig at a 99% identity threshold, each of the three assemblies recovers more than 87.3% of the reference, with MEGAHIT recovering the most – 89.3% of the reference (Table 7).

**Table 7:** Contig accuracy measured by reference coverage with strict alignment.

| Assembly | % covered |
| --- | --- |
| MEGAHIT | 89.3% |
| IDBA | 87.7% |
| MetaSPAdes | 83.4% |

### Individual genome statistics vary widely in the assemblies

We computed the NGA50 for each individual genome and assembly in order to compare assembler performance on genome recovery (see left panel of Figure 2). The NGA50 statistics for individual genomes vary widely, but there are consistent assembler-specific trends: IDBA yields the lowest NGA50 for 28 of the 51 genomes, while MetaSPAdes yields the highest NGA50 for 32 of the 51 genomes.

**Figure 2:**
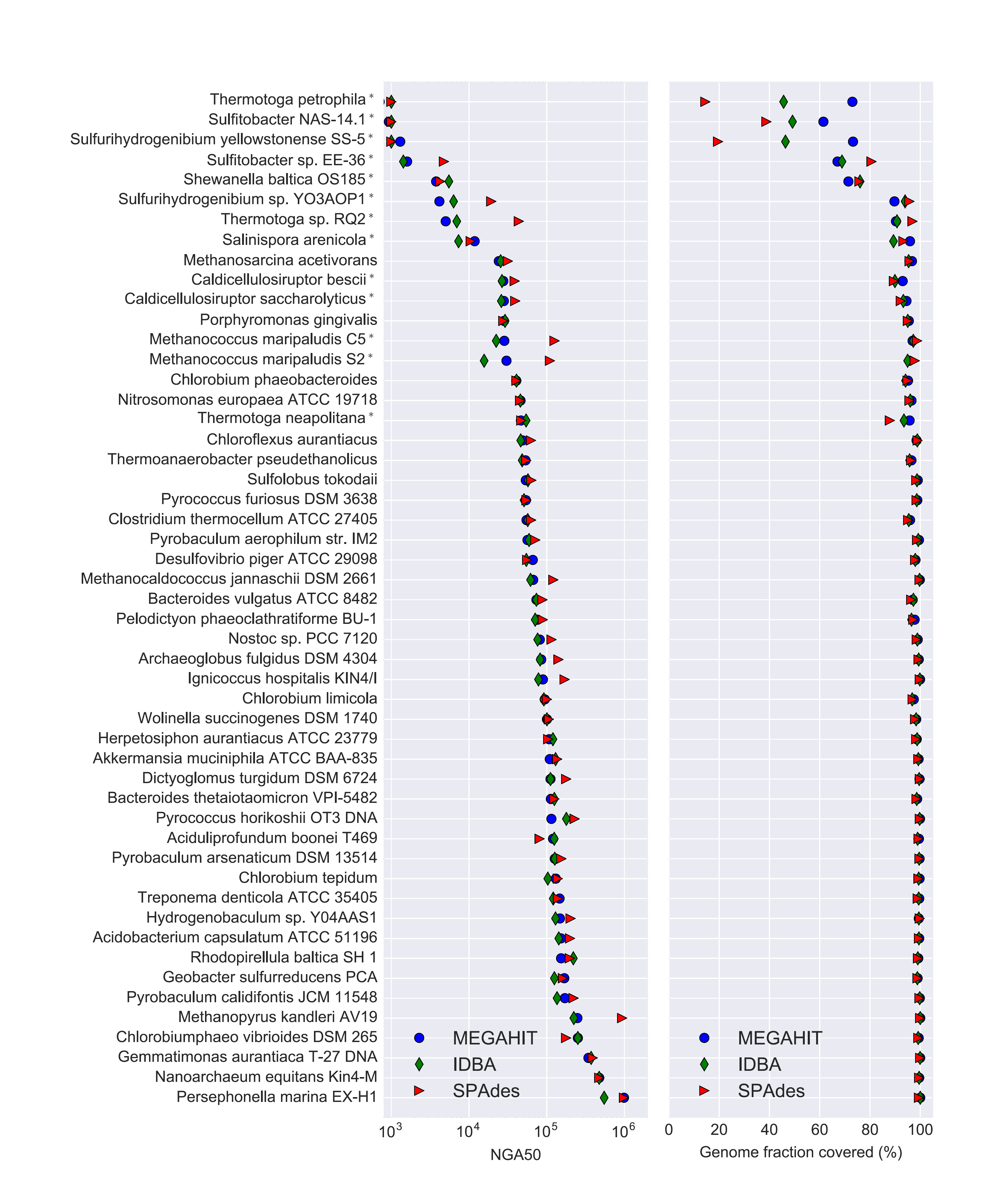
NGA50 and genome fraction covered, by genome and assembler. A ’*’ after the name indicates the presence of at least one other genome with > 2% Jaccard similarity at k=31 in the community. Where NGA50 cannot be calculated due to poor coverage, a marker is placed at 1kb.

We also evaluated aligned coverage per genome for each of the three assemblies (right panel, Figure 2). We found that 13 of the 51 genomes were missing 5% or more of bases in at least one assembly, despite all 51 genomes having 99% or higher read- and 51-mer coverage.

There are 12 genomes with k=31 Jaccard similarity greater than 2% to other genomes in the community, and these (denoted by ’*’ after the name) typically had lower NGA50 and aligned coverage numbers than other genomes. In particular, these constituted 12 of the 13 genomes missing 5% or more of their content, and the lowest eight NGA50 numbers.

### Longer contigs are less likely to be chimeric

Chimerism is the formation of contigs that include sequence from multiple genomes. We evaluated the rate of chimerism in contigs at three different contig length cutoffs: 500bp, 5kb, and 50kb (Table 8). We found that the percentage of contigs that match to the genomes of two or more different species drop as the minimum contig size increases, to the point where only the MEGAHIT assembly had a single chimeric contig longer than 50kb. Overall, chimeric misassemblies were rare, with no assembler generating more than 30 chimeric contigs out of thousands of total contigs.

**Table 8:** Chimeric contigs by contig length.

| Assembly | > 50kb | > 5kb | > 500 bp |
| --- | --- | --- | --- |
| IDBA | 0 | 1 | 7 (0.06%) |
| MEGAHIT | 1 | 4 | 14 (0.13%) |
| MetaSPAdes | 0 | 3 | 30 (0.48%) |

### The unmapped reads contain strain variants of reference genomes

Approximately 4.8 million reads (4.4%) from the QC data set did not map anywhere in the reference provided by the authors of [12]. We extracted and assembled these reads in isolation using MEGAHIT, yielding 6.5 Mbp of assembly in 1711 contigs > 500bp in length. We then did a k-mer inclusion analysis of this assembly against all of the GenBank genomes at k=31, and estimated the fraction of the k-mers that belonged to different species (Table 9). We find that 51.1% of the k-mer content of these contigs positively match to a genome present in GenBank but not in the reference metagenome.

**Table 9:** GenBank genomes detected in assembly of unmapped reads

| match | GenBank genome |
| --- | --- |
| 44.1% | *Fusobacterium* sp. OBRC1 |
| 23.0% | *P. ruminis* strain ML2 |
| 18.2% | *Thermus thermophilus* HB8 |
| 7.7% | *P. ruminis* strain CGMCC |
| 8.2% | *Enterococcus faecalis* M7 |
| 7.3% | *F. nucleatum* 13_3C |
| 3.7% | *F. nucleatum* subsp. *polymorphum* |
| 2.9% | *Fusobacterium hwasookii* |
| 1.0% | *E. coli* isolate YS |
| 1.7% | *F. nucleatum* subsp. *polymorphum*, alt. |
| 1.9% | *F. nucleatum* subsp. *vincentii* |

To verify these assignments, we aligned the MEGAHIT assembly of unmapped reads to the GenBank genomes in Table 9 with NUCmer using “loose” alignment criteria. We found that 1.78 Mbp of the contigs aligned at 99% identity or better to these GenBank genomes. We also confirmed that, as expected, there are no matches in this assembly to the full updated reference metagenome.

We note that all but the two *P. ruminis* matches and the *E. coli* isolate YS are strain variants of species that are part of the defined community but are not completely present in the reads (see Table 2). For *Proteiniclasticum ruminis,* there is no closely related species in the mock community design, and very little of the MEGAHIT assembly aligns to known *P. ruminis* genomes at 99%. However, there are many alignments to *P. ruminis* at 94% or higher, for approximately 2.73 Mbp total. This suggests that the unmapped reads contain at least some data from a novel species of *Proteiniclasticum*; this matches the observation in [12] of a contaminating genome from an unknown *Clostridium* spp., as at the time there was no *P. ruminis* genome.

## Discussion

### Assembly recovers basic content sensitively and accurately

All three assemblers performed well in assembling contigs from the content that was fully present in reads and k-mers. After length filtering, all three assemblies contained more than 95% of the reference (Table 6); even with removal of secondary alignments, more than 87% was recovered by each assembler (Table 7). About half the constituent genomes had an NGA50 of 50kb or higher (Figure 2), which, while low for current Illumina single-genome sequencing, is sufficient to recover operon-level relationships for many genes.

### The presence of multiple closely related genomes confounds assembly

In agreement with CAMI, we also find that the presence of closely related genomes in the metagenome causes loss of assembly [3]. This is clearly shown by Figure 2, where 12 of the bottom 14 genomes by NGA50 (left panel) also exhibit poor genome recovery by assembly (right panel). Interestingly, different assemblers handle this quite differently, with e.g. MetaSPAdes failing to recover essentially any of *Thermotoga petrophila,* while MEGAHIT recovers 73%. The presence of nearby genomes is an almost perfect predictor that one or more assembler will fail to recover 5% or more - of the 13/51 genomes for which less than 95% is recovered, 12 of them have close genomes in the community. Interestingly, very little similarity is needed - all genomes with Jaccard similarity of 2% or higher at k=31 exhibit these problems.

The *Shewanella baltica* OS185 genome is a good example: there are two strain variants, OS185 and OS223, present in the defined community. Both are present at more than 99% in the reads, and more than 98% in 51-mers, but only 75% of *S. baltica* OS185 and 50% of *S. baltica* OS223 are recovered by assemblers. This is a clear case of “strain confusion” where the assemblers simply fail to output contigs for a substantial portion of the two genomes.

Another interest of this study was to examine cross-species chimeric assembly, in which a single contig is formed from multiple genomes. In Table 8, we show that there is relatively little cross-species chimerism. Surprisingly, what little is present is length-dependent: longer contigs are less likely to be chimeric. This might well be due to the same “strain confusion” effect as above, where contigs that share paths in the assembly graphs are broken in twain.

### MEGAHIT performs best by several metrics

MEGAHIT is clearly the most efficient computationally, outperforming both MetaSPAdes and IDBA in memory and time (Table 4). The MEGAHIT assembly also included more of the reads than either IDBA or MetaSPAdes, and omitted only 0.4% more of the unique 51-mers from the reads than IDBA. MEGAHIT covered more of the reference genome with both loose and strict alignments (Table 6 and Table 7), with little duplication. This is clearly because of MEGAHIT's generally superior performance in recovering the genomes of closely related strains (Figure 2, right panel). The sum “fraction of genome recovered” is arguably the most important measure of a metagenome assembler (see [5] in particular) and here MEGAHIT excels for individual genomes even in the presence of strain variation.

In general other studies have found that MEGAHIT excels in recovery of sequence through assembly [3, 16] and is considerably more computationally efficient than most other assemblers [3, 15]. However, studies have also shown that MEGAHIT produces more misassemblies than other assemblers [3] and performs poorly on high coverage portions of the data set [5] Thus while we can recommend MEGAHIT as a good first assembler, we can also not unambiguously recommend it as the only assembler to use.

When comparing details of sequence recovery between the assemblers, the assembly content differs by only a small amount when loose alignments are allowed: all three assemblers miss more content (approximately 2.5% of the reference) than they generate uniquely (1.7% or less). In addition to preferring no one assembler over any other, this suggests that combining assemblies may have little value in terms of recovering additional metagenome content. The genome alignment statistics in Figure 2 suggest that much of this differential assembly content is due to the impact of strains.

### The missing reference may be present in strain variants of the intended species

Several individual genomes are missing in measurable portion from the QC reads (Table 2), and many QC reads (4.4% of 108m) did not map to the full reference metagenome. These appear to be related issues: upon analysis of the unmapped reads against GenBank, we find that many of the contigs assembled from the unmapped reads can be assigned to strain variants of the species in the mock community (Table 9) and align closely to the identified genomes. This suggests that the constructors of the mock community may have unintentionally included strain variants of *Fusobacterium nucleatum, Thermus thermophilus* HB27, and *Enterococcus faecalis;* note that the microbes used were sourced from the community rather than the ATCC (M. Podar, pers. communication). In addition, we detect what may be portions of a novel member of the *Proteiniclasticum* genus in the assembly of these reads - this is likely the *Clostridium* spp. detected through amplicon sequencing in [12].

Without returning to the original DNA samples, it is impossible to conclusively confirm that unintended strains were used in the construction of the mock community. In particular, our analysis is dependent on the genomes in GenBank: the genomes we detect in the contigs are clearly closely related to GenBank genomes not in the reference metagenome, based on k-mer analysis and contig alignment. However, GenBank is unlikely to contain the exact genomes of the actually included strain variants, rendering conclusive identification impossible.

## Conclusions

Overall, assembly of this mock community works well, with good recovery of known genomic sequence for the majority of genomes. All three assemblers that we evaluated recover similar amounts of most genomic sequence, but (recapitulating several other studies [3, 5, 15]) MEGAHIT is computationally the most efficient of the three. We note that assembly resolves substantial portions of several previously undetected strain variants, as well as recovering a substantial portion of a novel *Proteiniclasticum* spp. that was detected via amplicon analysis in [12], suggesting that assembly is a useful complement to amplicon or reference-based analyses.

The presence of closely related strains is a major confounder of metagenome assembly, and causes assemblers to drop considerable portions of genomes that (based on read mapping and k-mer inclusion) are clearly present. In this relatively simple community, this strain confusion is present but does not dominate the assembly. However, real microbial communities are likely to have many closely related strains and any resulting loss of assembly would be hard to detect in the absence of good reference genomes. While high polymorphism rates in e.g. animal genomes are known to cause duplication or loss of assembly, some solutions have emerged that make use of assumptions of uniform coverage and diploidy [31]. These solutions cannot however be transferred directly to metagenomes, which have unknown abundance distributions and strain content.

An additional concern is that metagenome assemblies are often performed after pooling data sets to increase coverage (e.g. [4, 32]); this pooled data is more likely to contain multiple strains, which would then in turn adversely affect assembly of strains. This may not be resolvable within the current paradigm of assembly, which focuses on outputting linear assemblies that cannot properly represent strain variation. The human genomics community is moving towards using *reference graphs,* which can represent multiple incompatible variants in a single data structure [33]; this approach, however, requires high-quality isolate reference genomes, which are generally unavailable for environmental microbes.

Long read sequencing (and related technologies) will undoubtedly help resolve strain variation in the future, but even with highly accurate long-read sequencing, current sequencing depth is still too low to resolve deep environmental metagenomes [34, 35]. It is unclear how well long error- prone reads (such as those output by Pacific Biosciences SMRT [36] and Oxford Nanopore instruments [37]) will perform on complex metagenomes: with high error rates, deep coverage of each individual genome is required to achieve accurate assembly, and this may not be easily obtainable for complex communities. Single-molecule barcoding (e.g. 10X Genomics [38]) and HiC approaches [39] show promise but these remain untested on well-defined complex communities and are still challenged by the complexity of complex environmental metagenomes; see [40, 41, 42].

Much of our analysis above depends on having a high-quality “mock” metagenome. While computationally constructed synthetic communities and computational “spike-ins” to real data sets can provide valuable controls (e.g. see [15] and [43]) we strongly believe that standardized communities constructed *in vitro* and sequenced with the latest technologies are critical to the evaluation of both canonical and emerging tools, e.g. efforts such as [44]. From the perspective of tool evaluation, we must disagree somewhat with Vollmers et al. [5]: good metagenome tool evaluation necessarily depends on mock communities that are as realistic as we can make them. Likewise, from the perspective of bench biologists, actually sequencing real DNA is critical because it can evaluate confounding effects such as kit contamination [45]. Large-scale studies of computational approaches systematically applied to mock communities such as CAMI [3] can then provide fair comparisons of entire toolchains (wet + dry) applied to these mock communities.

We omitted two important questions in this study: binning and choice of parameters. We chose not to evaluate genome binning because most binning strategies either operate post-assembly (see e.g. [46]), in which case the challenges with assembly discussed above will apply; or require multiple samples (e.g. [47]), which we do not have. We also chose to use only default parameters with all three assemblers, for two reasons. First, we are not aware of any effective automated approaches for determining the “best” set of parameters or evaluating the output for metagenome assemblers, other than those integrated into the assemblers themselves (e.g. the choice of k-mer sizes by MEGAHIT and MetaSPAdes), and absent such guidance we do not feel comfortable blessing any particular set of parameters; here the choice of default parameters is parsimonious (and also see [48] for the dangers of poorly chosen objective functions). Second, any parameter exploration pipeline would not only need to be automated but would need to run multiple assemblies, whose time and resource usage should be measured; in this case, any comparison based on runtime of the parameter choice pipeline should naturally favor MEGAHIT because of its advantage in computational efficiency.

## Author contributions

SA, LI and CTB developed, tested, and executed the analytical pipeline. SA and CTB created the tables and figures and wrote the paper.

## Acknowledgments

We thank Michael R. Crusoe and Phillip T. Brooks for input on analysis and pipeline development. We thank Migun Shakya, Mircea Podar, Jiarong Guo, Harald R. Gruber-Vodicka, Juliane Wippler, Krista Ternus, and Stephen Turner for valuable comments on drafts of this manuscript.

